# Interactive Effects of Venlafaxine and Thermal Stress on Zebrafish (*Danio rerio*) Inflammatory and Heat Shock Responses

**DOI:** 10.1101/2022.11.18.517121

**Authors:** A.V. Weber, B.F. Firth, I. G. Cadonic, P.M. Craig

## Abstract

Venlafaxine (VFX), a commonly prescribed antidepressant often detected in wastewater effluent, and acute temperature elevations from climate change and increased urbanization, are two environmental stressors currently placing freshwater ecosystems at risk. This study focused on understanding if exposure to VFX impacts the agitation temperature (T_ag_) and critical thermal maximum (CT_max_) of zebrafish (*Danio rerio*). Additionally, we examined the interactive effects of VFX and acute thermal stress on zebrafish heat shock and inflammatory immune responses. A 96 hour 1.0 μg/L VFX exposure experiment was conducted, followed by assessment of thermal tolerance via CT_max_ challenge. Heat shock proteins and pro-inflammatory immune cytokines were quantified through gene expression analysis by quantitative PCR (qPCR) on *hsp 70, hsp 90, hsp 47, il-8, tnfα*, and *il-1β* within gill and liver tissue. No significant changes in agitation temperature between control and exposed fish were observed, nor were there any differences in CT_max_ based on treatment. Unsurprisingly, *hsp 47, 70, and 90* were all upregulated in groups exposed solely to CT_max_, while only *hsp 47* within gill tissue showed signs of interactive effects, which was significantly decreased in fish exposed to both VFX and CT_max_. No induction of an inflammatory response occurred. This study demonstrated that environmentally relevant concentrations of VFX have no impact on thermal tolerance performance in zebrafish. However, VFX is capable of causing diminished function of protective heat shock mechanisms, which could be detrimental to freshwater fish populations and aquatic ecosystems as temperature spikes become more frequent from climate change and urbanization near watersheds.

**Summary Statement:** This study predicts the effects that climate change and anthropogenic pollutants may have on fish ability to tolerate elevated temperatures, and examines the physiologic challenges these stressors may introduce.

## Introduction

With the world population close to surpassing 8 billion, the impact of anthropogenic stress on aquatic environments is a growing concern. Climate and land use changes are increasing water temperature (Mohajerani et al., 2017; Poesch et al., 2016; Seneviratne et al., 2014; Somers et al., 2013; Stillman, 2019), and pollutant discharges introduce additional stressors to these environments (Daughton and Ternes, 1999). These factors culminate in adverse impacts on freshwater species, particularly those that that reside in relatively shallow habitats that are more susceptible to temperature changes because of lower thermal inertia (Morash et al., 2021).

Humans are responsible for the introduction of pharmaceutical and personal care products (PPCPs) into freshwater ecosystems (Arlos et al., 2015; Daughton and Ternes, 1999; Metcalfe et al., 2010). PPCPs are a specific class of environmental contaminant that can induce physiological effects in exposed organisms (Ebele et al., 2017). Venlafaxine (VFX), a known PPCP, is a commonly prescribed serotonin and norepinephrine reuptake inhibitor (SNRI) often detected downstream of wastewater treatment plants (WWTPs; (Metcalfe et al., 2010). The medication and its active metabolite, O-desmethyl venlafaxine (O-VFX), enter the effluent through human excretion, are released into freshwater ecosystems through WWTP outfall, and persist in aquatic environments at lower doses of its original prescribed form (Metcalfe et al., 2010). VFX and O-VFX are of particular interest as several studies have discovered the two to be more abundant than any other antidepressant in wastewater effluent (Gauvreau et al., 2022; Hodgson et al., 2020; Metcalfe et al., 2010) and previous work has demonstrated numerous adverse physiologic effects in exposed species, including decreases in embryo production (Galus et al., 2013), disruptions in metabolic responses to secondary stressors (Best et al., 2014), decreases in miRNA expression (Ikert and Craig, 2020), generational impacts on epigenetic responses (Luu et al., 2021), and reduced survival (Schultz et al., 2011).

Environmental contaminants can activate an immune response in exposed animals, however the influence of VFX specifically is less understood. Preliminary pilot experiments performed in our lab found a 96-hour 1.0 μ/L VFX exposure to zebrafish resulted in increased expression of pro-inflammatory cytokines, *il-8, tnfα, il-1β*. (Dawe and Craig, unpublished observations), which help to initiate and direct inflammatory immune responses. (Semple and Dixon, 2020; Uribe et al., 2011). Meagre (*Argyrosomus regius*) housed in VFX-spiked water resulted in promotion of cellular damage in exposed fish, suggesting VFX may possess the potential to initiate an inflammatory immune response (Maulvault et al., 2019). Conversely, VFX has also been reported to have anti-inflammatory properties, as recent findings have shown several anti-depressant medications are capable of depressing inflammatory action in mammals (Hajhashemi et al., 2015; Tynan et al., 2012). Thus, the impact of VFX on fish inflammatory responses remains unclear, highlighting a need for further study. Here, Il-1*β* (interleukin 1*β*), and Tnf*α* (tumor necrosis factor *α*,), proinflammatory cytokines responsible for initiating inflammatory responses, were analyzed along with Il-8 (interleukin 8), a chemokine that functions to recruit neutrophils and basophils to the site of infection and has been deemed a marker of immune system activation (Semple & Dixon, 2020).

Additionally, several studies have investigated the effects of aquatic contaminants on the heat shock response in fish, a protective stress mechanism that results in upregulation of heat shock proteins (Hsp). Hsp’s act as molecular chaperones, aiding in protein folding and preventing or reversing any misfolding that may occur following exposure to stressors like heat, contaminants, and pathogens (Abdel-Gawad and Khalil, 2013; Lindquist and Craig, 1988; Mitra et al., 2018; Roberts et al., 2010). In this study, we examined the effects of effluent exposure on the expression of three Hsp’s of interest: Hsp 47, 70, and 90, all of which have previously shown to be affected by contaminant exposure. One study on catfish (*Rita rita*) found that both Hsp 70 and Hsp 47 were significantly upregulated in fish sampled from polluted environments compared to their non-polluted counterparts (Mitra et al., 2018). Similarly, Hsp 90 increased expression in response to a variety of environmental pollutants across several fish species, such as common carp (*Cyprinus carpio*) and ornate wrasse (*Thalassoma pavo*; (Xing et al., 2015; Zizza et al., 2017). Though common contaminants in aquatic environments are capable of inducing upregulation of Hsp’s, the effect of VFX alone on Hsp expression has not been examined in fish. A study on rat cell culture found that exposure to 10 μM VFX initially increased expression of Hsp 70, proposing that VFX may be capable of regulating HSP expression, however this effect has yet to be analyzed in fish (Yu et al., 2010).

In addition to contaminants, the occurrence of extreme heat waves is becoming more common (Stillman, 2019). These heat waves, in tandem with rainwater runoff from urban heat islands, can result in acute temperature spikes in freshwater ecosystems (Mohajerani et al., 2017; Somers et al., 2013; Stillman, 2019). Temperature increases pose a serious threat to poikilothermic organisms (such as fish) that are incapable of maintaining their internal body temperature, potentially harming physiologic processes (Deutsch et al., 2008; Ficke et al., 2007; Hochachka and Somero, 2002). The inflammatory immune response is known to be directly modulated by temperature, with some levels capable of enhancing immune function (Alcorn et al., 2002; Le Morvan et al., 1997), but supra-optimal levels can diminish appropriate responses (Dominguez et al., 2004). Elevated temperature can also indirectly impact immune function, as production of cortisol during stress has immunosuppressive effects (Cortés et al., 2013; Petrovsky, 2001; Tort, 2011). Specific pro-inflammatory cytokines (Tnfα, Il-1β, and Il-8) have previously shown enhanced expression following temperature elevations; however, considering the immune system appears to have an optimal temperature range, it is unclear how an acute heat spike may impact cytokine expression in zebrafish (Perezcasanova et al., 2008; Polinski et al., 2013; Singh et al., 2008).

Temperature increases also induce initiation of the heat shock response. Hsp 47, 70, and 90 have all been reported to be capable of upregulation following acute heat stress (Fangue et al., 2011; Manzon et al., 2022; Zhang et al., 2014). Fangue et al. determined that Hsp 70 was inducible in situations of acute thermal stress introduced through Critical Thermal Maximum (CT_max_), a method of temperature ramping used to determine an organism’s thermal tolerance. CT_max_ can provide insight into how contaminant-exposed fish are able to overcome acute changes in temperature, and was used in this study to gain a better idea on how VFX exposure may influence fish ability to tolerate additional environmental stressors. Prior exposure experiments have indicated that pollutants like pesticides can reduce thermal tolerance (Op de Beeck et al., 2017), yet others have shown that fish from effluent polluted environments had no changes in thermal tolerance compared to non-polluted sites (Nikel et al., 2021). Although no studies have investigated VFX impacts on CT_max_, WWTP effluent can increase oxygen consumption which could potentially manifest as limitations in handling higher temperatures (Mehdi et al., 2018). Drawing inspiration from Fangue et al. (2011), CT_max_ was utilized in this project to assess for thermal tolerance and also as a method of inducing acute thermal stress to investigate the inflammatory and heat shock response after VFX exposure.

Environmental stressors are rarely experienced individually, rather, combinations of stressors can result in exacerbated effects of pollutants, specifically in the presence of elevated temperature. Thermal stress increases metabolic rate in fishes, increasing ventilation and ultimately increasing exposure levels to toxic contaminants (Cairns et al., 1975; Maulvault et al., 2018). This is especially true for effluent-derived contaminants considering that WWTP effluent has been shown to cause thermal enhancement near discharge sites (Environment Canada, 2001; Mehdi et al., 2019). Furthermore, since most WWTPs are located in urban centers, warmed rainwater runoff due to the heat island effect can lead to acute temperature spikes in effluent polluted waters (Mohajerani et al., 2017; Somers et al., 2013). Climate change will exacerbate this effect further due to increased frequency and severity of heat waves (Seneviratne et al., 2014; Stillman, 2019). Thus, knowing how temperature spikes will interact with contaminants is imperative for aquatic species. It is therefore necessary that multiple interacting stressors be studied in order to accurately represent aquatic environments and gain a better understanding of the dangers these interactions may pose to aquatic ecosystems. This study attempts to understand if VFX exposure impacts zebrafish thermal tolerance and investigates if exposure to both acute thermal stress and VFX have interactive effects on the heat shock and inflammatory immune responses of zebrafish. We hypothesized that VFX-exposed fish would have decreased ability to tolerate higher temperatures, and predicted that fish exposed to both VFX and thermal stress would have increased levels of heat shock proteins and inflammatory cytokines.

## Materials and Methods

### Experimental Design

Adult zebrafish of mixed sex were obtained from a local supplier (Big Al’s Kitchener, Ontario, Canada) and held in a recirculating Z-HAB system (Pentair Aquatic Eco-Systems Inc., Apopka, Florida, USA). Tanks were maintained at 27° C, pH 7.5, and conductivity of ∼670 μS, under 12-hour light dark cycles. All experiments performed in this study were in accordance with the Canadian Council of Animal Care guidelines as reviewed by the University of Waterloo Animal Care Committee (AUP #40989).

An acute 96-hour VFX exposure experiment was designed with two treatment groups: an exposed group spiked with 1.0 μg/L of VFX and a non-exposed control group (Figure 1). Zebrafish of mixed sex were transferred to a 15 L glass tank with three tank replicates per treatment. Tanks were oxygenated via air stone, continuously filtered, and held at 27° C for fish to acclimate for 3 days prior to exposure. At the start of the exposure, aquarium filters were removed and VFX (Millipore-Sigma-Aldrich, Oakville, Ontario, Canada) was introduced into the exposed group tanks at a concentration of 1.0 μg/L. This concentration was chosen because VFX levels in effluent exposed waters can reach 1.0 μg/L and has been previously shown to impact transcript levels in fish (Gauvreau et al., 2022; Luu et al., 2021; Metcalfe et al., 2010). Throughout the experiment, fish were fed Gemma 300 (Skretting, Westbrook, Maine, USA) to satiety once daily followed by a 50% water change 1 hour after feeding to minimize nitrogenous waste build-up. Following daily water changes, re-addition of VFX was dosed to maintain an overall concentration of 1.0 μg/L. Water quality parameters (pH, Nitrite, Nitrate, and Ammonia) were checked 1 hour after VFX dosage using a Freshwater Master Test Kit (API, Chalfont, Pennsylvania, USA). Upon completion of the 96 hours, thermal tolerance was measured in a subset of fish from each tank. The remaining fish were euthanized using buffered 0.5 g/L MS-222 and sampled to serve as a non-heat exposed baseline. Length, weight, and sex of each fish were recorded, and gill and liver tissue were dissected, immediately frozen on dry ice, and stored in individual cryotubes at -80° C until further use. Based on this study design, 4 experimental groups were established: control/no CT_max_ fish referred to as control baseline, control/CT_max_ fish referred to as control-heat, VFX/no CT_max_ fish referred to as VFX baseline, and VFX/CT_max_ fish referred to as VFX-heat (Figure 1).

**Figure 1:**
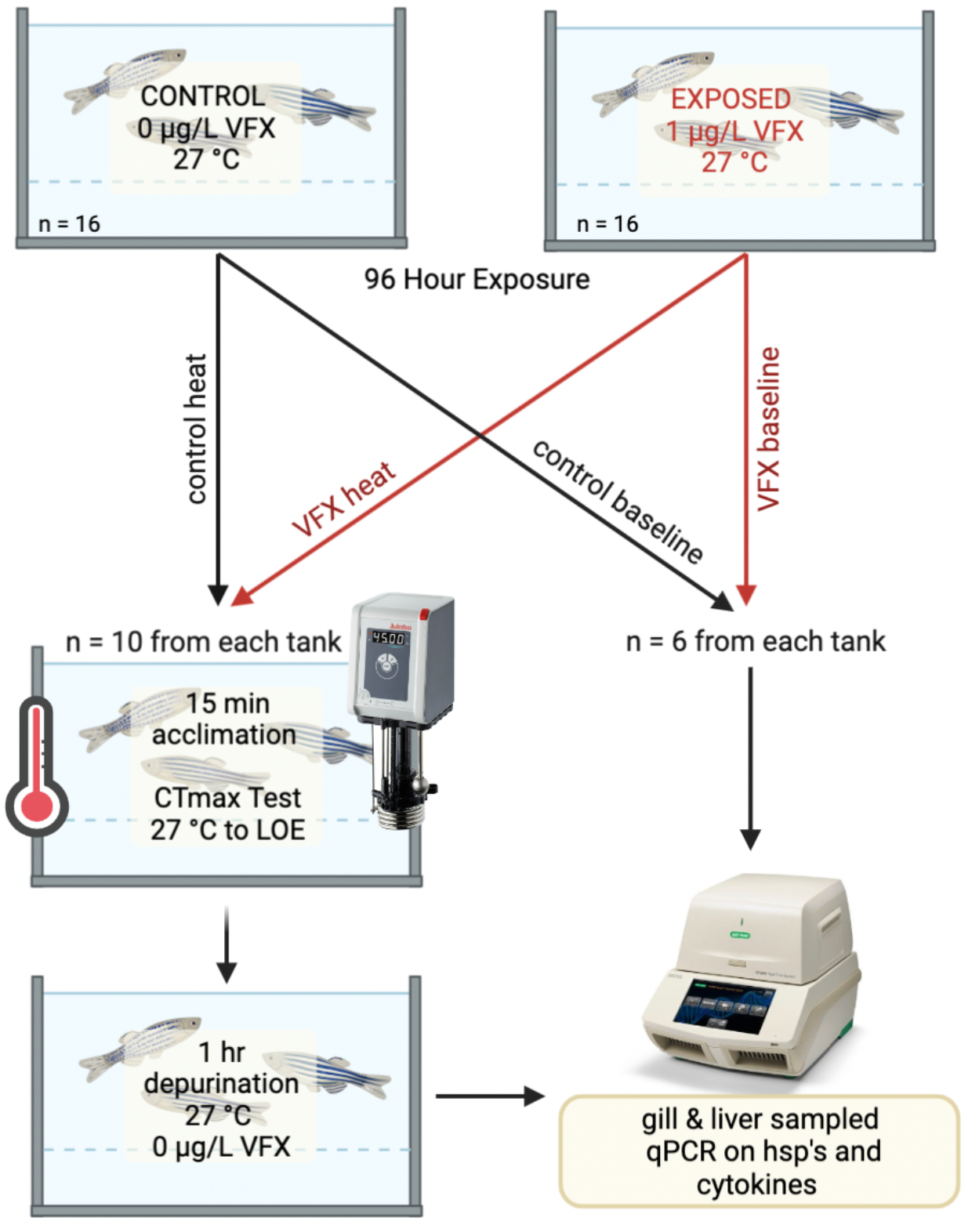
Graphical representation of experimental design. A 96-hour exposure experiment was conducted with two treatment groups: a 1.0 μg/L VFX exposed group and a non-exposed control. Upon completion of the 96 hours, thermal tolerance was measured in the control heat and VFX heat experimental groups. The remaining fish were euthanized to serve as a control baseline and VFX baseline. All groups had liver and gill tissue sampled and expression of hsp’s and pro-inflammatory cytokines was determined via qPCR.

### VFX Quantification

Twice over the course of the 96 hours, water samples were taken from each treatment tank to confirm VFX concentrations and lack of VFX in control tanks. Samples were collected at least 1 hour after daily VFX re-spiking and stored in amber glass bottles at -20 ° C until analysis. Analysis of samples was conducted according to the protocol described in Luu et al. (2021). VFX was quantified through solid phase extraction (SPE) in Oasis HLB cartridges (6cc, 500 mg, Waters Corporation, Milliford, Massachusetts, USA) followed by liquid chromatography and tandem mass spectrometry using a Sciex API 32000 QTRAP LC-MS/MS system (ABSciex; Concord, Ontario, Canada). The method detection limit (MDL) in a 500 mL samples is 1/ng/L. 100 mL samples were extracted in this experiment, so the detection limit was calculated to be 5 ng/L based on the original MDL.

### Assessment of Thermal Tolerance

Once the 96-hour VFX exposure had completed, zebrafish from each treatment underwent an assessment of thermal tolerance determined via CT_max_. Fish were fasted 24 hours prior to the CT_max_ procedure. For the CT_max_ trials, 5 zebrafish were transferred to a breeder box placed within a 30 L glass aquarium and left to acclimate for 15 minutes at 27°C. After acclimation, temperature was increased by 0.33°C per minute using a Julabo portable immersion circulator (Julabo, Seelbach, Baden-Würrtemberg, Germany). Fish behavior was visually monitored, specifically examining for agitation temperature (T_ag_) and loss of equilibrium (LOE). Agitation temperature was marked as the temperature at which fish appear to become distressed and begin to take on more erratic swimming behaviors in an attempt to search for cooler waters (McDonnell and Chapman, 2015). LOE is the point at which thermal tolerance is reached; at this temperature fish cannot maintain their position in the water column and go belly up (Becker and Genoway, 1979). Upon observation of agitation behavior and LOE, temperature of occurrence was recorded. Fish that had reached their thermal maximum were placed into a holding tank of 27°C, 0.0 μg /L VFX for a 1-hour depurination period to allow time to mount a heat shock response, as adapted from the methods used in Fangue et al., 2011. Euthanasia and sampling of gills and liver was conducted using the same method utilized on baseline fish. No mortalities occurred due to the CT_max_ procedure. All trials were recorded via GoPro and re-watched to confirm agitation and LOE temperatures. Experimental film was analyzed blindly, through randomized sorting and renaming of video files performed by a simple Python script. The percent of fish displaying signs of temperature agitation, and percent reaching LOE were recorded at specific temperature points as it increased incrementally. Temperature was log transformed and plotted with either the percentages of agitation or LOE as a dose response curve to determine the EC50 value for each treatment (Figure 3A).

### Tissue Analysis

Liver and gill tissue RNA were extracted following Craig et al. (2013). Per 100 mg of tissue, 1 mL of Trizol (Sigma-Aldrich, Oakville, Ontario, Canada) was added and homogenized by an OMNI TH handheld tissue homogenizer (Kennesaw, Georgia, USA). Chloroform was added to the Trizol-tissue solution and samples were centrifuged to separate the two liquid layers. Supernatants were removed and precipitated with 100% isopropyl alcohol. Again, samples were centrifuged. The remaining precipitate was washed and centrifuged with 75% EtOH twice. EtOH was removed and samples were pulse spun and air dried to ensure the pellet had no trace EtOH. Samples were reconstituted in water. RNA concentration was quantified using a SpectraMax 190 (San Jose, California, USA) and 500 ng of RNA was used for cDNA synthesis, using a Qiagen QuantiTect Reverse Transcription kit (Hilden, North-Rhine Westphalia, Germany). cDNA samples were stored at -20° C until subsequent analysis. The relative expression levels of the genes: *hsp 70, hsp 90, hsp 47, il-8, tnfα, il-1β*, (Table 1) were determined using quantitative PCR on a BioRad CFX96 Touch Thermal Cycler (Hercules, California, USA) with a sample size of n = 6 fish per treatment derived from one tank exposure replicate (n = 5 for VFX baseline in liver tissue due to RNA concentration being too low). Each reaction contained 2 μL of diluted cDNA, 5 μL of BioRad SYBR Green Master Mix, 1 μL of forward and reverse primers, and 1 μL of water. cDNA was incubated at 95° C for 30 s, denatured at 95° C for 10 s, and annealed at 60° C for 20 s. Fluorescence was then detected and followed by 39 more denaturation and annealing cycles. Gene expression was normalized to several housekeeping genes: *β-actin* and ribosomal protein subunit (*rps 18*) for gill samples, and *β-actin* and *ef1a* for liver, all of which were deemed stable between treatment groups (Table 2).

**Table 1:**
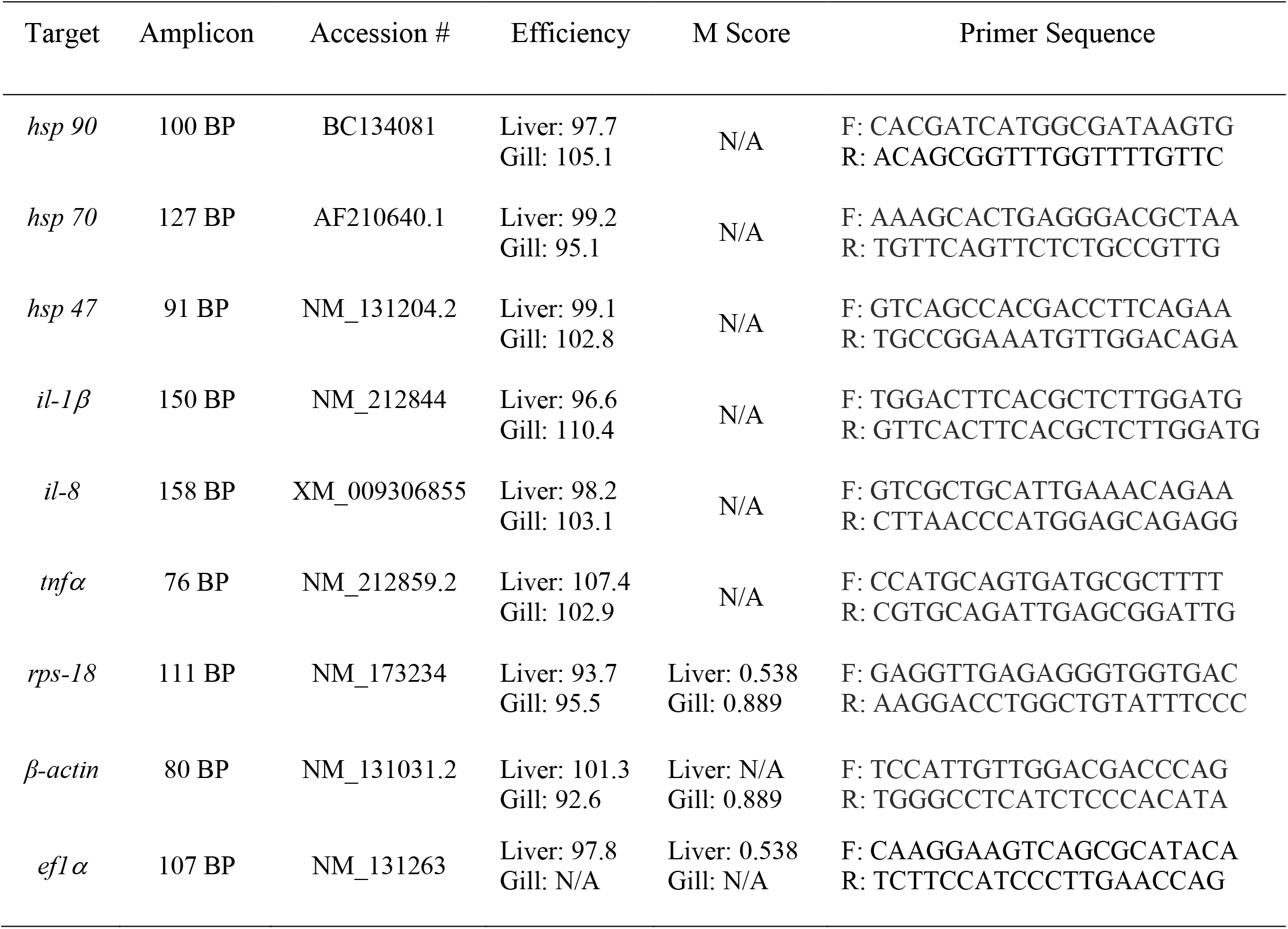
mRNA primers of genes of interest used for RT-qPCR. All primer sequences are listed in the 5’ to 3’ direction with F representing forward and R reverse primers.

### Statistical Analysis

All statistical analyses were performed by GraphPad Prism 8.1.2 using a p value cutoff of 0.05 (GraphPad, San Diego). An independent t-test was completed to assess for any significant differences in the CT_max_ values of VFX-exposed and non-exposed. Dose response curves were created for both agitation temperature and LOE with incremental temperature increases as dosage and percent agitated or percent LOE as response. EC50 values were determined and used in an independent samples t-test to compare for any significant differences in 50% agitation temperature and 50% LOE between exposed and non-exposed treatment groups. For all CT_max_ data, individual data points were averaged within each experimental replicate and analyses were conducted between replicate averages. Hsp’s and pro-inflammatory cytokine gene expression data were log-transformed to pass normality testing for parametric statistics (gill *hsp 90* expression was normally distributed and therefore was not log-transformed). Significant changes in expression were determined via Two-Way Analysis of Variance (ANOVA) followed by a Tukey post hoc test to analyze for interactive effects of VFX exposure and acute thermal stress.

## Results

### Water Quality

Venlafaxine quantification analyses ran on control and VFX exposed water samples confirmed control tanks to have an average of 0.0 μg/L of VFX and exposure tanks 1.076 ± 0.020 μg/L, with values presented as mean ± SEM.

### Agitation Temperature and Critical Thermal Maximum

Thermal tolerance was not significantly different (t-test, t_4_ = 0.8489, P = 0.4438) between VFX-exposed (41.25° C ± 0.13) and non-exposed (40.93° C ± 0.36) treatment groups (Figure 2). VFX exposed fish (EC50 34.00° C) showed signs of temperature agitation 0.68 ° C earlier than control fish (EC50 34.68° C). Conversely, LOE was reached 0.20° C earlier in control fish (EC50 41.02° C) than VFX-exposed (EC50 41.22° C). No significant differences were seen between control and VFX EC50 values in either agitation temperature (t-test, t_4_ = 1.251, p = 0.2790) or LOE (t-test, t_4_ = 1.294, p = 0.2652; Figure 3B).

**Figure 2:**
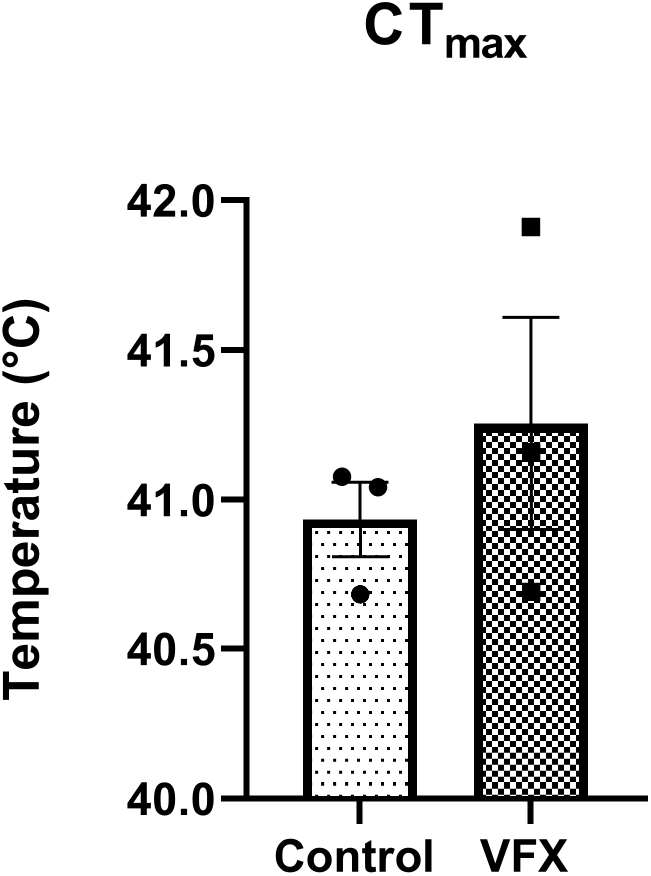
Critical thermal maximum of zebrafish from control and VFX-exposed treatment groups. Different letters denote significant differences between groups (p < 0.05; n = 3. Individual data points are shown for each exposure replicate CT_max_ average and bars represent mean ± standard error.

**Figure 3:**
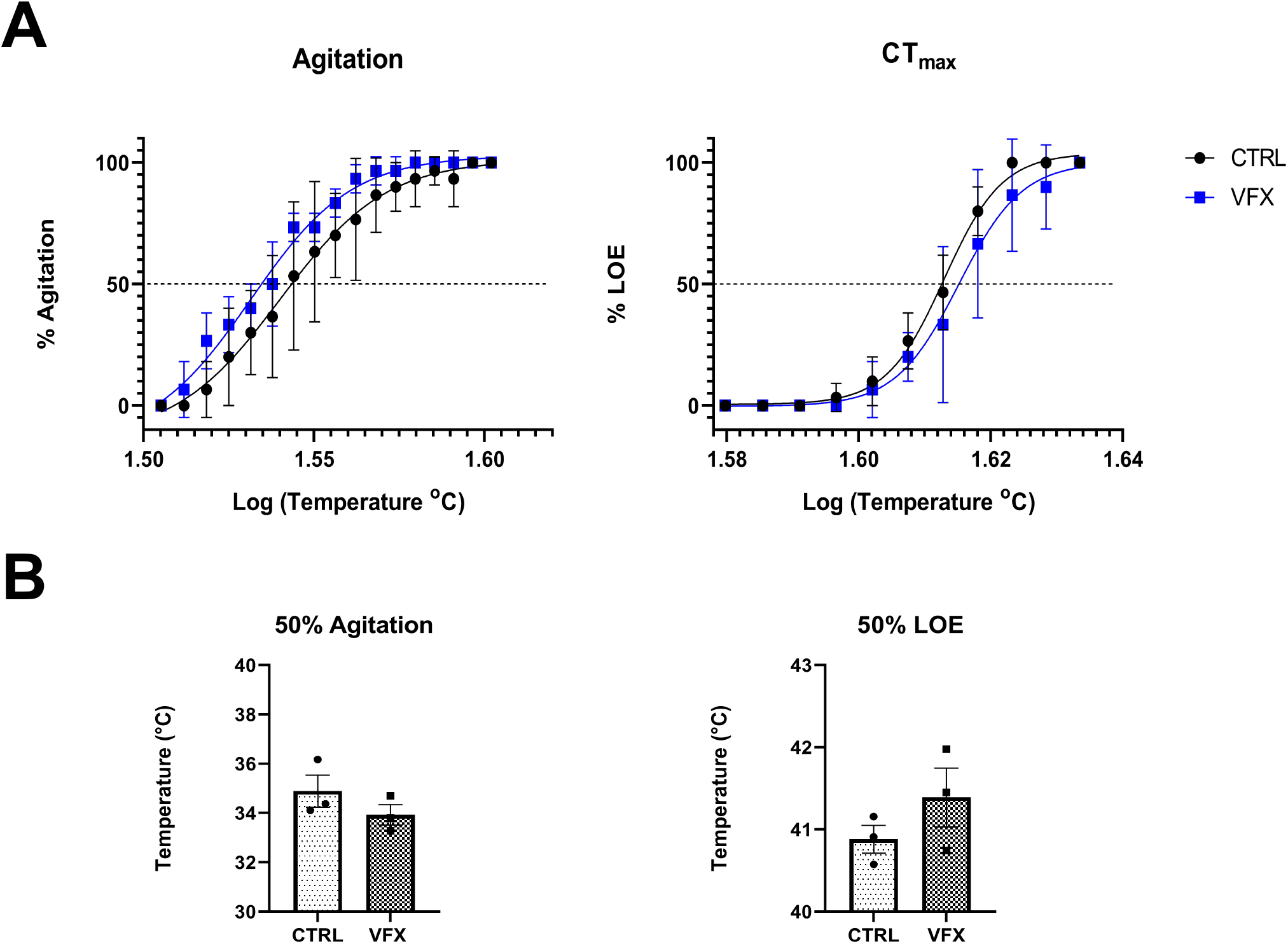
Percent of fish displaying signs of agitation and reaching LOE at specific temperature increments. Recorded following a blind review of the CT_max_ experiment on film, temperature points were log transformed and plotted a dose response curve in order to determine the point of 50% agitation and 50% LOE for each treatment (A) (n = 3 exposure replicates). Temperature agitation, control fish: EC50 34.68° C, VFX exposed fish: EC50 34.00° C. LOE control fish: EC50 41.02° C, VFX exposed fish: EC50 41.22° C. Dots represent temperature points at which percentages were recorded. EC50 values of agitation temperature and LOE were analyzed using an independent samples t-test (B) (p < 0.05, n = 3 exposure replicates).

### RT-qPCR Gene Expression Analysis

All pro-inflammatory cytokine expression in liver tissue was not significantly different between treatment group (Two-Way ANOVA, p > 0.05), heat exposure (Two-Way ANOVA, p > 0.05), or the interaction term between treatment group and heat exposure (Two-Way ANOVA, p > 0.05; Table S1; Figure 4A). Similarly in gill tissue, pro-inflammatory cytokine expression was not significantly different between treatment group (Two-Way ANOVA, p > 0.05), heat exposure (Two-Way ANOVA, p > 0.05) or the interaction term between treatment group and heat exposure (Two-Way ANOVA, p > 0.05; Table S2; Figure 4B).

**Figure 4:**
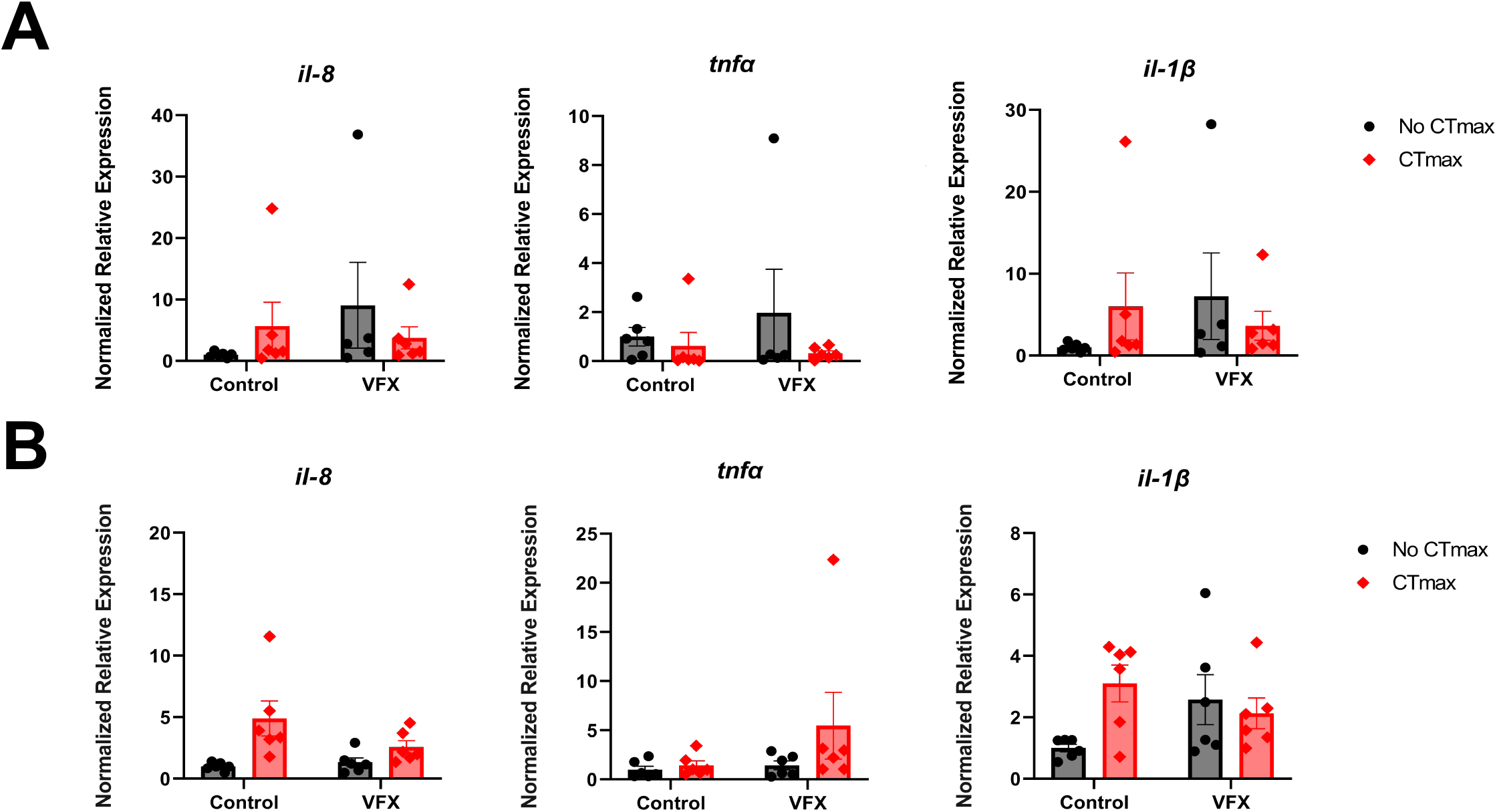
Pro-inflammatory cytokine gene expression. The relative gene expression (mean ± SEM) of *il-8, tnfα*, and *il-1β*, in zebrafish liver (A) and gill (B) in either control or VFX-exposed fish followed by either presence or absence of CT_max_ thermal challenge. Bars with differing symbols show significant differences between them as determined by a Two-Way ANOVA with Tukey’s post hoc test (p < 0.05; n = 6). Dots represent individual data points. Gene expression figures are presented as untransformed data for clarity.

Within liver tissue all hsp’s were significantly upregulated in zebrafish that underwent a CT_max_ thermal challenge, regardless of exposure group (Figure 5A). *Hsp 70* was significantly different between heat exposure (Two-Way ANOVA, F_1,19_ = 317.7, p = < 0.0001) while treatment group (Two-Way ANOVA, F_1,19_ = 0.007899, p = 0.9301) and the interaction between treatment group and heat exposure was not significantly different (Two-Way ANOVA, F_1,19_ = 0.1702, p = 0.6845; Figure 5A). *Hsp 70* elicited the greatest upregulation out of all examined hsp’s, increasing by about 2,000 fold between control baseline and control-heat fish, and then roughly 1,000 fold VFX baseline and VFX-heat fish (Tukey HSD, p < 0.0001; p < 0.001). Likewise, *hsp 90* was significantly different between heat exposure (Two-Way ANOVA, F_1,19_ = 276.4, p = < 0.0001). Treatment group (Two-Way ANOVA, F_1,19_ = 0.2212, p = 0.6435) and the interaction between treatment group and heat exposure were not significantly different (Two-Way ANOVA, F_1,19_ = 0.006506, p= 0.9366; Figure 5A). Expression of *hsp 90* increased 500 fold in control-heat and 300 fold in VFX-heat exposed fish (Tukey HSD, p < 0.0001; p < 0.001). *Hsp 47* also was significantly different between heat exposure (Two-Way ANOVA, F_1,19_ = 59.3, p = < 0.0001) with no significant differences observed between treatment group (Two-Way ANOVA, F_1,19_ = 0.2109, p = 0.6513) and the interaction between treatment group and heat exposure (Two-Way ANOVA, F_1,19_ = 1.891, p = 0.1851; Figure 5A). *Hsp 47* appeared to have a more minor, but still significant increase with expression increasing more than 20 fold following thermal challenge in control group fish and 8 fold in the VFX-heat treatment group (Tukey HSD; p < 0.0001; p = 0.0017 respectively).

**Figure 5:**
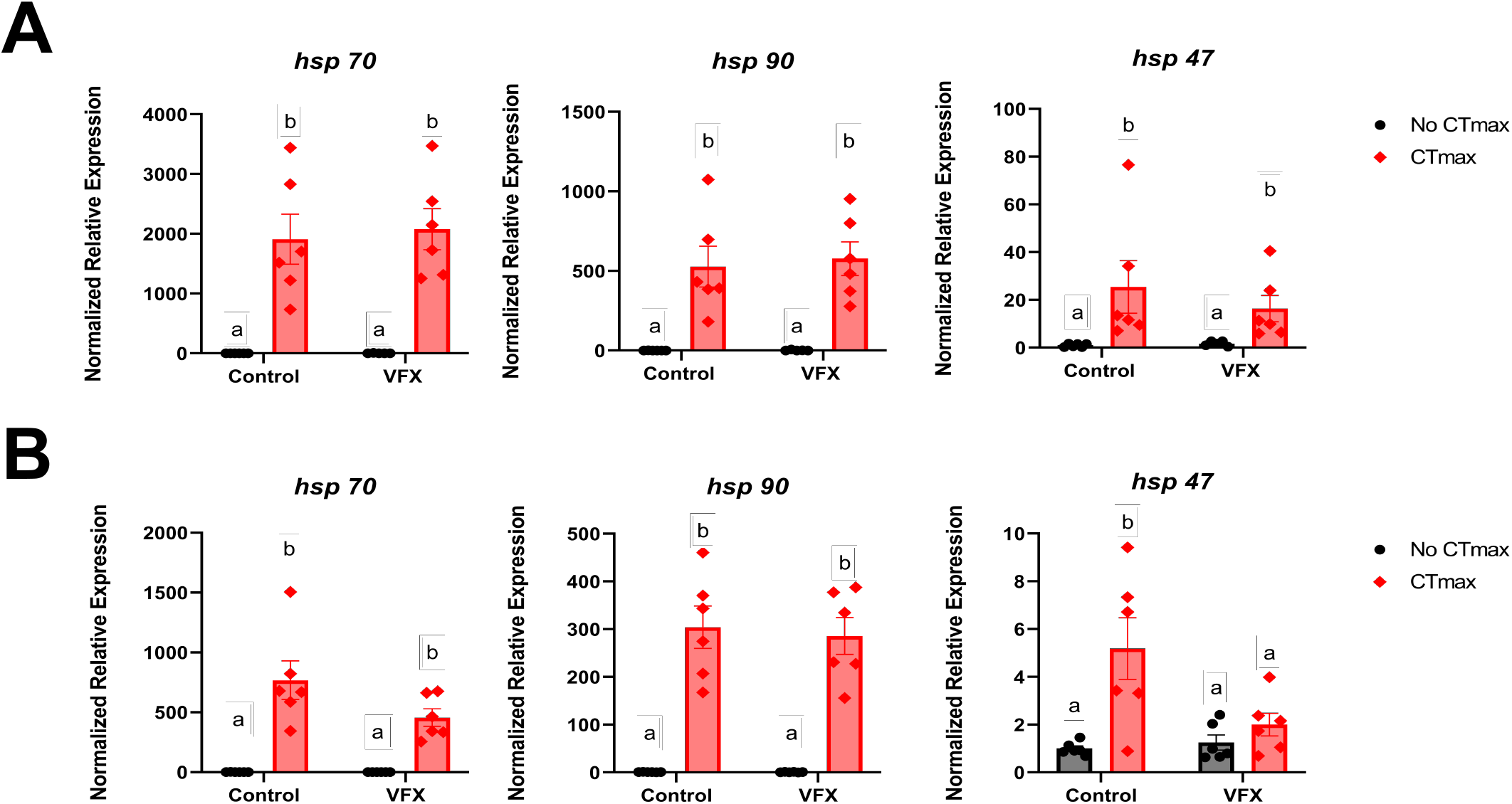
Heat shock protein gene expression. The relative gene expression (mean ± SEM) of *hsp 70, hsp 90*, and *hsp 47* in zebrafish liver (A) and gill (B) in either control or VFX-exposed fish followed by either presence or absence of CT_max_ thermal challenge. Bars with differing symbols show significant differences between them as determined by a Two-Way ANOVA with Tukey’s post hoc test (p < 0.05; n = 6). Dots represent individual data points. Gene expression figures are presented as untransformed data for clarity.

Similar patterns were noted in gill *hsp 70* and *hsp 90* expression; both significantly increased in all fish exposed to CT_max_ (Two-Way ANOVA, F_1,20_ = 8.559, p = 0.0084; Two-Way ANOVA, F_1,20_ = 99.93, p = < 0.0001 respectively; Figure 5B). Furthermore, both the treatment group (Two-Way ANOVA, F_1,20_ = 2.215, p = 0.1523; Two-Way ANOVA, F_1,20_ = 0.09774, p = 0.7578 respectively) and the interaction between treatment group and heat exposure were not significantly different in neither *hsp 70* or *hsp 90* (Two-Way ANOVA, F_1,20_ = 2.215, p = 0.1523; Two-Way ANOVA, F_1,20_ = 0.09573, p = 0.7602 respectively; Figure 5B). Expression of *hsp 70* in gill tissue increased ∼800 fold in control-heat and ∼700 fold in VFX-heat fish (Tukey HSD, p < 0.001; p < 0.001). Expression of *hsp 90* saw ∼250 fold increase in fish exposed to control-heat and ∼300 fold increase in VFX-heat fish (Tukey HSD, p < 0.001; p < 0.001). It is noteworthy that *hsp 47* was the only gene of interest examined in the study that showed interactive effects between acute heat stress and VFX exposure (Two-Way ANOVA, F_1,20_ = 4.984, p = 0.0372). Expression of *hsp 47* was significantly reduced by approximately 60% in VFX-heat compared to control-heat fish (Tukey HSD, p = 0.0293). Within control fish, expression was upregulated roughly 4 fold in control heat fish (Tukey HSD, p = 0.0119); in VFX-exposed fish, no significant differences were observed between VFX baseline and VFX-heat (Tukey HSD, p = 0.9891). Overall, for both gill and liver tissue, exposure to VFX alone did not induce a heat shock response in any of the hsp’s.

## Discussion

This study aimed to determine if VFX exposure impacts fish thermal tolerance and alters the heat shock and inflammatory immune responses to acute thermal stress in zebrafish. We demonstrated that a 96-hour exposure to an environmentally relevant concentration of VFX is not sufficient in decreasing zebrafish thermal tolerance. Neither VFX, CT_max_, nor interaction of the two, were found to have significant impacts on expression of pro-inflammatory cytokines. However, CT_max_ was capable of inducing upregulation of all hsp transcripts measured and it was found that VFX dampened the heat shock responses, as seen as lowered *hsp 47* expression levels in gill tissue.

### 1.0 Thermal Tolerance

VFX exposure did not alter thermal tolerance as no significant differences were seen between the temperatures at which agitation and LOE occurred between VFX-exposed and non-exposed fish (Figures 2 & 3). It is possible that the environmentally relevant concentration of 1.0 ug/L or the 96-hour exposure period simply are not enough to introduce significant physiological changes that would present adverse effects on fish thermal tolerance. Our findings here coincide with similar studies that assessed the impacts of contaminated environmental sites on CT_max_ performance (Jayasundara et al., 2017; Nikel et al., 2021). Though both Jayasundara et al. (2017) and Nikel et al. (2021), observed other changes caused by exposure to contaminants, including alterations in body size, mitochondrial function and metabolic processes, no changes in CT_max_ between exposed and non-exposed treatment groups occurred in either study. Thus, while VFX may be capable of inducing alternate physiological changes (Best et al., 2014; Ikert and Craig, 2020; Mehdi et al., 2019), it can be concluded that these effects are not sufficient to result in decreased ability to tolerate warming waters.

One challenge of the assessment of thermal tolerance was determining the points of agitation temperature. Considering that zebrafish are a pelagic species and often swim erratically under normal, non-stressed conditions, pinpointing the onset of agitation proved to be difficult. A blind assessment of each trial was utilized to mitigate bias; however, it is still possible that the assessment of agitation performed in this study may not be representative of when individual fish experienced thermally-induced agitation. Comparison of agitation temperature between exposure groups was of particular interest considering that it is a behavioral measure that could be impacted in fish exposed to an antidepressant medication with known neurological effects (Gauvreau et al., 2022). Numerous studies have implied VFX’s role in behavioral changes like predation behavior and escape responses in fish, so additional studies on VFX-induced behavioral impacts specifically relating to temperature tolerance is worth further investigation (Bisesi et al., 2014; Painter et al., 2009). Although no significant conclusions were drawn here, studying a benthic species would be advantageous in effectively elucidating the impact of VFX exposure on agitation temperature.

### 2.0 Inflammatory Response

It is unlikely that VFX and acute heat stress have any serious inflammatory impacts on zebrafish considering that no changes were observed in pro-inflammatory cytokine expression in both gill and liver tissue across all treatment groups. Surprisingly, VFX alone presented no inflammatory effects. Previous studies have shown that environmentally relevant concentrations of other PPCPs, specifically (non-steroidal anti-inflammatory drugs) NSAIDs and ibuprofen were capable of inducing inflammatory responses and increasing cytokine production within exposed fish (Hoeger et al., 2005; Zhang et al., 2021). Likewise, 1.0 μg/L VFX exposure in various darter species (*Etheostoma* spp.) caused significant upregulation of *il-6* and *caspase 9*, pro-inflammatory and apoptotic markers, in the gill (Dawe et al., under review). However, these differences were relatively subdued (2-fold) compared to a true inflammatory response. An active immune response would be expected to result in cytokine upregulations of more than 100 fold (Commins et al., 2010; Zou and Secombes, 2016). Considering that no significant changes in pro-inflammatory cytokine expression occurred in the current study following VFX exposure, and the recent darter studies found only 2-fold increases, it appears that the environmentally relevant concentration of 1.0 ug/L VFX alone is not sufficient to induce an active inflammatory response in zebrafish.

All immune transcripts (*il-1β, il-8, tnfα)* were unaffected by the acute temperature stress (Figure 4). However, previous studies have shown that acute heat stress can induce short term immune enhancement (Tort, 2011). It is particularly interesting that the pro-inflammatory cytokines within this study did not follow this trend. It is plausible that the lack of significance in upregulation observed in heat groups may be due to the 1-hour depurination period that followed CT_max_. Cytokine responses to acute stressors are known to have a shorter half-lives compared to those activated in response to chronic stressors (Tort, 2011). Therefore, cytokine expression could have returned to control levels by the time sampling occurred. Further, heat has been shown to enhance the inflammatory immune response, often having synergistic effects when experienced in addition to an inflammatory stimulant like lipopolysaccharide (LPS; (Polinski et al., 2013; Tort, 2011). Considering that VFX had no impact on the immune transcripts measured, it is not surprising that the introduction of acute heat shock, or the interaction between heat and VFX, caused no additional upregulation of cytokines. To better understand impact of VFX, temperature, and the interactive effects of the two on inflammation, a live pathogen challenge would be a more effective approach to elucidate how these stressors may be influencing the inflammatory immune response.

### 3.0 Heat Shock Response

While numerous studies have investigated the impacts of aquatic contaminants on Hsp expression, few have illustrated the effect of effluent-derived contaminants, specifically VFX. As expected, Hsp expression increased drastically in both liver and gill after acute temperature increases, affirming similar results presented in Fangue et al (2011), that deemed CT_max_ capable of inducing a heat shock response. Examination of the interactive effects of VFX exposure and acute thermal stress found that VFX may diminish the heat shock response, as *hsp 47* within gill tissue was unaffected by CT_max_ (Figure 5). This result is concerning considering that the gill is indirect contact with the environment and has important roles in regulating oxygen transport; a VFX-induced dampened heat shock response could introduce harmful physiologic effects relating to complications in oxygen uptake, such as decreased oxygen transport ability and thus reduced oxygen delivery to tissues. It is unclear why only *hsp 47*, localized only within the endoplasmic reticulum and functioning specifically in collagen folding and assembly, showed interactive effects and not *hsp 90* or *70*, which are implicated in more general protein folding and expressed ubiquitously (Iwama et al., 1999; Zhang et al., 2014). Furthermore, the physiological implications of this dampened expression should be investigated further to identify if VFX impacts collagen folding after heat stress. Previous work has suggested that there are different regulatory mechanisms behind *hsp 70* and *47*, however the specifics underlying these differences and why one chaperone might be impacted by VFX exposure and not the other remain unknown (Lele et al., 1997). Similarly, VFX’s method of dampening hsp expression in this study is unclear. Yu et al (2010) has postulated that VFX may be capable of causing translocation of glucocorticoid receptors into the nucleus which could result in inhibition of heat shock factor (HSF), a transcription factor that controls expression of heat shock proteins, ultimately preventing the formation of an appropriate heat shock response. However, this provides no insight into why expression of some hsp’s and not others may be impacted by VFX. Studies utilizing different concentrations of VFX could provide insight into whether this antidepressant modulates heat shock responses. As well, the reasoning behind why *hsp 47* expression is sensitive to VFX is puzzling and requires further investigation.

### 4.0 Conclusions

Collectively, this study demonstrated that environmentally relevant concentrations of VFX have no impact on thermal tolerance performance in zebrafish. Furthermore, VFX, and interaction between VFX and acute thermal stress present no induction of an inflammatory immune response. VFX is capable of causing diminished function of protective heat shock mechanisms which could prove to be detrimental to freshwater fish populations and aquatic ecosystems as temperature spikes become more frequent from climate change and increased urbanization near watersheds.

## Acknowledgments

The authors would like to thank Leslie Bragg and the Servos Lab for assistance with VFX quantification, and Neil Brubacher for creation of the Python script used in blind video analysis. Experimental design graphic was created with BioRender.com. This research was funded through a Natural Sciences and Engineering Research Council (NSERC) Discovery Grant to PMC.

